# *Lrat-Cre* Exhibits Widespread Expression Beyond Hepatic Stellate Cells Across Multiple Tissues

**DOI:** 10.64898/2026.01.09.698660

**Authors:** Kyungchan Kim, Solaema Taleb, Jisun So, Jamie Wann, Hyun Cheol Roh

## Abstract

Hepatic stellate cells (HSCs) play a central role in liver fibrosis, shifting from quiescent vitamin A-storing cells to activated, myofibroblast-like cells that secrete collagen and other profibrotic factors^1^. HSCs have thus become a major focus in liver fibrosis research, and several Cre driver lines have been created to target HSCs in mice. However, early Cre lines had significant limitations. Glial fibrillary acidic protein (*Gfap*)*-Cre* labels only a subset of HSCs and also induces recombination in cholangiocytes^2^. Collagen type I alpha 1 (*Col1a1*)*-Cre* and alpha-smooth muscle actin (*αSMA*)*-Cre/CreERT2* primarily label activated myofibroblasts and broadly mark portal fibroblasts and vascular smooth muscle cells^3,4^. Platelet-derived growth factor receptor beta (*Pdgfrβ*)*-Cre* reliably labels HSCs but also recombines pericytes and smooth muscle cells, limiting its specificity^5^. The introduction of lecithin-retinol acyltransferase (*Lrat*)*-Cre* marked a major advance, offering highly specific labeling of quiescent and activated HSCs and rapidly becoming the most widely used driver for HSC tracing and genetic perturbation^2^. However, the extrahepatic expression of *Lrat-Cre* remains incompletely understood. This is a critical limitation, given that liver biology is closely coordinated with other organs to maintain systemic metabolism. Addressing these gaps is essential for the accurate interpretation of HSC-specific genetic models in liver biology.

## Results

We crossed *Lrat-Cre* mice with the Nuclear tagging and Translating Ribosome Affinity Purification (NuTRAP) reporter line^6^, which enables Cre-inducible expression of ribosome-bound green fluorescent protein (GFP) and mCherry-labeled nuclei (Figure 1A). As expected, *Lrat-Cre*; NuTRAP mice exhibited GFP expression specifically in HSCs (Figure 1B and Supplemental Figure 1A). We next performed gene expression analysis using GFP-based ribosome pulldown to isolate HSC-specific mRNA alongside whole-liver RNA. Key HSC markers, such as *Lrat, Acta2*, and *Des*, were highly enriched by 60-to 200-fold relative to whole liver (Figure 1C). We further extended this analysis with RNA-seq, identifying 2,033 genes significantly enriched and 1,493 genes significantly depleted in HSCs (Figure 1D and Table S1). Pathway analysis confirmed strong enrichment of extracellular matrix organization pathways and depletion of lipid metabolic processes in HSCs (Figures 1E-F). Overall, these findings confirm that *Lrat-Cre* specifically marks HSCs within the liver, consistent with previous characterizations^2^.

**Figure 1.**
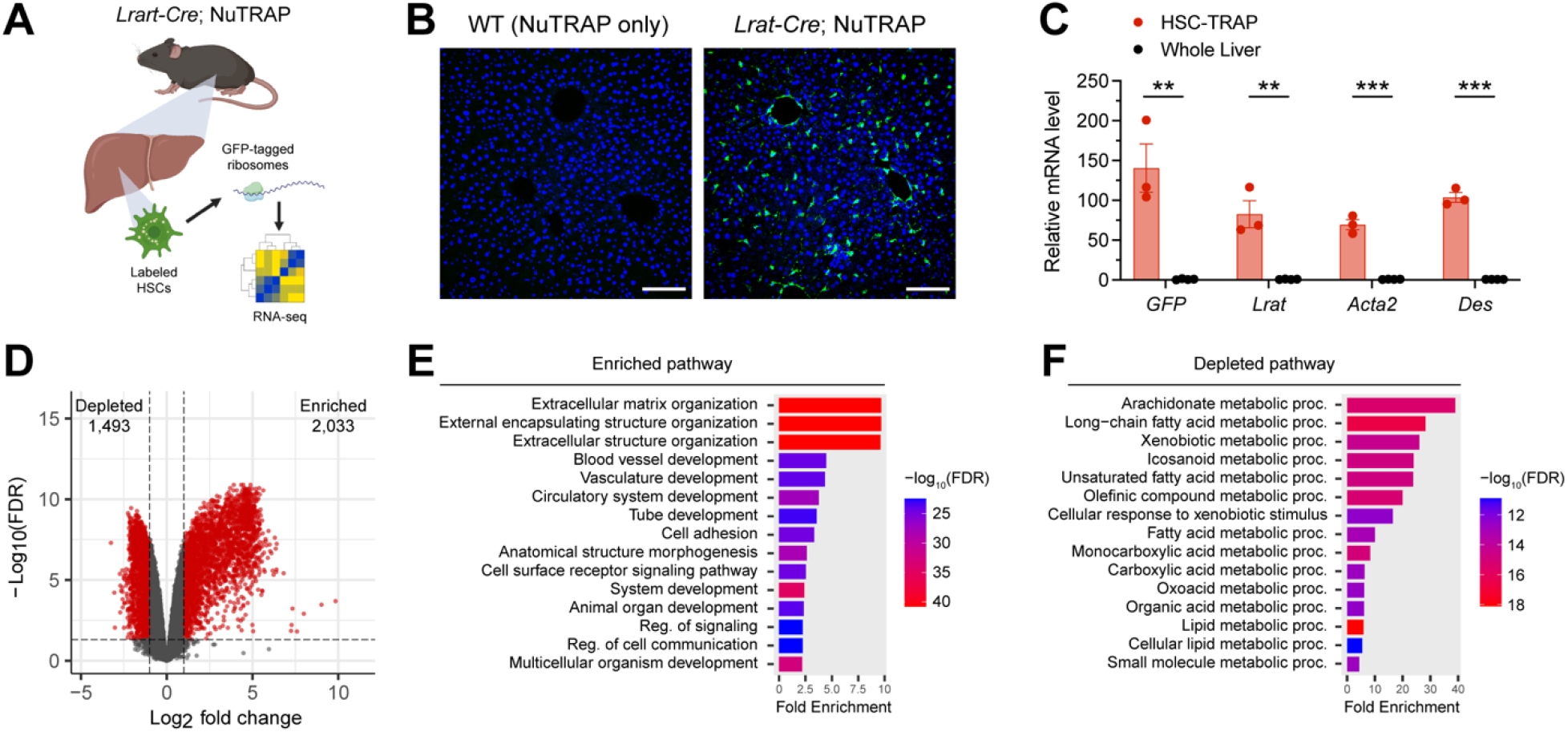
HSC-specific labeling and transcriptomics using *Lrat-Cre*; NuTRAP. (A) Schematic of the *Lrat-Cre*; NuTRAP approach. (B) Liver immunofluorescence showing GFP (green) and Hoechst (nuclei, blue). Scale bar: 100 µm. (C) qPCR analysis of HSC markers in HSC-TRAP versus whole liver RNA. Data shown as mean ± SEM (n=3). \*\**p* < 0.01; \*\*\**p* < 0.005. (D) Volcano plot of differentially expressed genes. (E-F) GO biological process analysis of enriched (E) and depleted (F) pathways in HSCs.

To examine *Lrat-Cre* activity outside the liver, we measured *GFP* and *Cre* mRNA expression across multiple tissues from *Lrat-Cre*; NuTRAP mice. *GFP* was essentially absent in NuTRAP-only wild-type (WT) controls, although a minor background signal was detected in a few tissues (Figure 2A), likely due to genomic DNA contamination. In *Lrat-Cre*; NuTRAP mice, *GFP* was nearly undetectable in spleen and pancreas, but all other tissues showed significant *GFP* expression, similar to or markedly higher than in the liver, with lung displaying nearly a 100-fold increase. Interestingly, *Cre* expression showed a different pattern (Figure 2B). The eye and testis displayed the highest *Cre* mRNA levels, consistent with the known strong endogenous *Lrat* expression in these tissues^7^. Most other GFP-positive tissues expressed *Cre* at 1-7-fold relative to liver (Figure 2B). However, *Cre* expression was generally lower than *GFP* expression, suggesting that these tissues contain cells with a history of Cre activity that activated *GFP* expression earlier, while ongoing *Cre* expression may be limited. Furthermore, immunofluorescence staining confirmed GFP-positive cells across multiple tissues (Figure 2C). We observed GFP in adipocytes and progenitors within adipose tissues (Figure 2C, Supplemental Figures 1B-C); in myotubes of skeletal muscle (Figure 2C); in enterocytes along the intestinal villi (Figure 2C, Supplemental Figure 2A); and in alveolar epithelial cells of the lung (Figure 2C, Supplemental Figure 2B). GFP-positive cells were particularly abundant in the intestine and lung (Figure 2C). Taken together, these findings demonstrate that *Lrat-Cre* is active and drives recombination in a wide range of non-hepatic tissues at a surprisingly high level.

**Figure 2.**
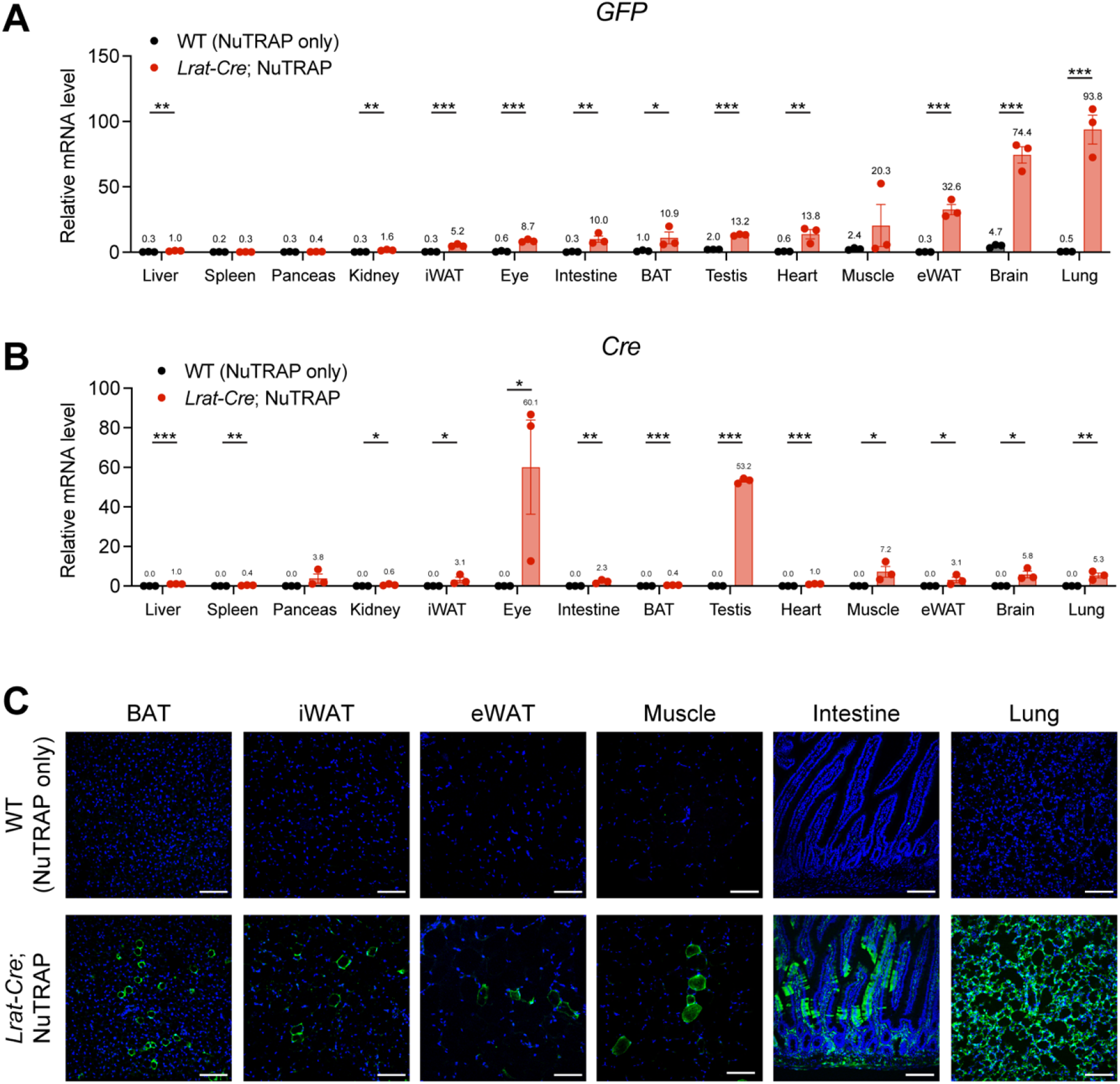
*Lrat-Cre* shows broad activity in non-hepatic tissues. (A-B) qPCR for *GFP* (A) and *Cre* (B) expression across tissues in WT and *Lrat-Cre*; NuTRAP mice. Data shown as mean ± SEM (n=3). \**p* < 0.05; \*\**p* < 0.01; \*\*\**p* < 0.005. (C) Immunofluorescence for GFP in the indicated tissues. Scale bars: 100 µm. BAT, brown adipose tissue; iWAT, inguinal white adipose tissue; eWAT, epididymal white adipose tissue.

Using NuTRAP reporter mice, we confirmed that *Lrat-Cre* specifically labels HSCs within the liver and generated *in vivo* HSC transcriptomic profiles that can serve as a useful reference for liver biology. Unexpectedly, however, we also observed substantial *Lrat-Cre* recombination across multiple non-hepatic tissues. While this may not affect its use for reporter-based HSC labeling, it becomes a major concern for genetic perturbation studies. In mouse models using *Lrat-Cre* to drive HSC-specific knockout or overexpression, recombination in metabolically important extrahepatic tissues, such as adipose tissue, skeletal muscle, heart, intestine, and brain, can significantly confound phenotypic interpretation. Liver phenotypes in these models may therefore arise from functional changes in these extrahepatic tissues rather than from HSCs themselves. For example, *Lrat-Cre*-mediated gene deletion or overexpression in the brain may influence feeding behavior; in skeletal muscle, physical activity or energy expenditure; and in the intestine, where we detected widespread recombination in enterocytes, nutrient absorption could be substantially altered. Thus, phenotypes observed in *Lrat-Cre*-based mouse models must be interpreted with caution. Orthogonal approaches, such as adeno-associated virus-mediated HSC-specific gene delivery or validation in primary HSC cultures, are essential to confirm HSC-intrinsic effects. Although inducible systems such as *Lrat-CreERT2*^8^ and *Lrat-rtTA*^9^ provide temporal control and may offer improved specificity, their extrahepatic activity has not yet been thoroughly characterized. In conclusion, our findings underscore the importance of carefully considering extrahepatic Cre activity when using *Lrat-Cre* for HSC-targeted genetic studies.

## Acknowledgments

We acknowledge the support provided by the Indiana University School of Medicine Center for Medical Genomics and the Center for Biological Microscopy.

## Abbreviations

BAT: (Brown adipose tissue),
Col1a1: (Collagen type I alpha 1)
eWAT: (Epididymal white adipose tissue)
GFAP: (Glial fibrillary acidic protein)
GFP: (Green fluorescent protein)
HSCs: (Hepatic stellate cells)
iWAT: (Inguinal white adipose tissue)
Lrat: (Lecithin-retinol acyltransferase)
NuTRAP: (Nuclear tagging and Translating Ribosome Affinity Purification
Pdgfrβ: (Platelet-derived growth factor receptor beta)
WT: (Wild-type)

