## Supplemental Information for "*Lrat-Cre* Exhibits Widespread Expression Beyond Hepatic Stellate Cells Across Multiple Tissues"

### MATERIALS AND METHODS

#### Animals

All animal procedures were approved by the Institutional Animal Care and Use Committee at the Indiana University School of Medicine. Mice were housed under a 12-hour light-dark cycle at 22 °C with ad libitum access to food and water. *Lrat-Cre* mice (Jackson Laboratory, 069595-JAX)[1] were crossed with NuTRAP reporter mice (Jackson Laboratory, 029899)[2] to generate *Lrat-Cre*; NuTRAP animals. Wild-type NuTRAP littermates lacking Cre served as controls. Male mice aged 4 months were used for all experiments, except where noted. For the high-fat, high-cholesterol, and cholate diet (HFCC; 60kcal% fat, 1.25% cholesterol, 1.25% cholate; Research Diets), 6-week-old mice were fed HFCC for 12 weeks.

#### Ribosome Pulldown for HSC-specific RNA Isolation

Ribosome pulldown was performed as previously described[3], with minor modifications. Frozen liver tissues were Dounce-homogenized on ice in homogenization buffer (50 mM Tris-HCl, pH 7.5; 12 mM $MgCl_2$; 100 mM KCl; 1% NP-40; 100 µg/mL cycloheximide; 1 mg/mL heparin; 2 mM DTT; RNasin; and protease inhibitors). Lysates were cleared by centrifugation (13,000 rpm, 10 min, 4 °C), and the resulting supernatant was incubated with anti-GFP antibody (5 µg/mL; Abcam, ab290) for 1 hour at 4 °C. Protein G Dynabeads prewashed in low-salt buffer (50 mM Tris-HCl, pH 7.5; 12 mM $MgCl_2$; 100 mM KCl; 1% NP-40; 100 µg/mL cycloheximide; 2 mM DTT) were added for 1 hour, followed by three washes in high-salt buffer (same as low-salt buffer but with 300 mM KCl). Bead-bound RNA was extracted using TRIzol (Thermo Fisher Scientific) according to the manufacturer’s instructions. For input RNA, 5% of the initial homogenate was mixed with TRIzol and processed in parallel to isolate whole liver RNA.

#### Quantitative Real-Time PCR (qPCR)

cDNA synthesis was performed with 500 ng RNA using the High-Capacity cDNA Reverse Transcription Kit (Applied Biosystems). qPCR was carried out using SYBR Green Master Mix (Applied Biosystems) on a QuantStudio 5 instrument. Gene expression was normalized to the housekeeping gene Rplp0, and fold change was calculated using the ΔΔCt method.

**Bulk RNA Sequencing (RNA-seq)**

RNA from HSC ribosome pulldown and whole liver were subjected to on-column DNase I treatment using the TURBO DNase Kit (Invitrogen). RNA quality was assessed with an Agilent Bioanalyzer. For library preparation, 100 ng of RNA underwent ribosomal RNA depletion using the NEBNext rRNA Depletion Kit (New England BioLabs), followed by first- and second-strand cDNA synthesis. Libraries were prepared using the Nextera XT DNA Library Preparation Kit (Illumina) and amplified by PCR. Indexed libraries were quantified by Qubit and analyzed on a Bioanalyzer for size distribution. Libraries were sequenced on an Illumina NovaSeq 6000.

**RNA-seq Data Analysis**

RNA-seq reads were quality-filtered and trimmed using fastp[4], then aligned to the mouse mm10 reference genome using STAR[5]. Gene-level counts were obtained with featureCounts[6]. Genes with low expression (counts per million [CPM] ≤ 1 in more than half of the samples) were removed prior to differential expression analysis using edgeR[7]. Differentially expressed genes were defined as those with average $\log_2$CPM ≥ 2, a $\log_2$ fold change ($\log_2$FC) ≥ 1, and a false discovery rate (FDR) < 0.05. Pathway enrichment analysis was performed using ShinyGO[8], with genes exhibiting $\log_2$FC ≥ 4 considered HSC-enriched and those with $\log_2$FC ≤ -1 considered HSC-depleted.

**Immunofluorescence Staining**

Processing, embedding, cryosectioning, and immunofluorescence staining were performed as previously described[9]. Briefly, tissues were fixed in 4% paraformaldehyde, cryoprotected in 30% sucrose, embedded in OCT, and sectioned at 15 μm (liver, muscle, and intestine), 30 μm (BAT and lung), or 50 μm (iWAT and eWAT). Sections were blocked in 5% donkey serum with 0.1% Triton X-100, incubated with primary antibodies overnight at 4 °C, washed, and incubated with fluorophore-conjugated secondary antibodies (1:300; Invitrogen), followed by Hoechst staining. Primary antibodies were used at the following final dilutions: goat anti-GFP (1:300; Novus, nb100-1678), rabbit anti-GFP (1:500; Abcam, ab290), rabbit anti-αSMA (1:100; Abcam, ab5694), goat anti-PLIN1 (1:100; Abcam, ab61682), goat anti-PDGFRα (1:50; R&D systems, AF1062), mouse anti-desmin (1:100, Dako, M0760), and rabbit anti-PLIN2 (1:100, Proteintech, 15294-1-AP). Images were acquired using an Olympus FV1000MPE confocal microscope from 3–5 sections per mouse (≥2 mice per group), and representative images are shown.

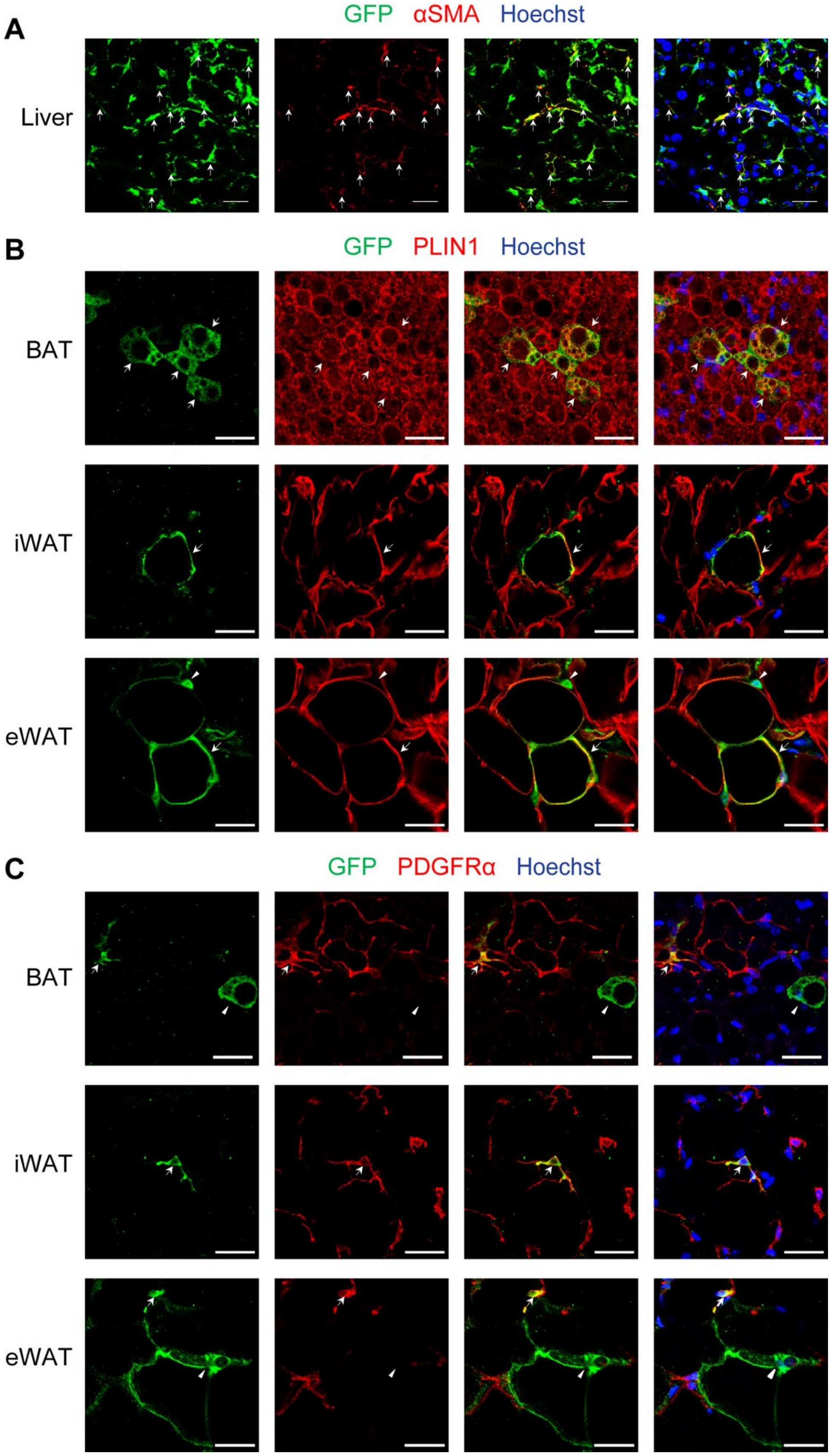
A
GFP αSMA Hoechst
Liver
B
GFP PLIN1 Hoechst
BAT
iWAT
eWAT
C
GFP PDGFRα Hoechst
BAT
iWAT
eWAT

**Supplemental Figure 1. Distribution of GFP-positive cells in *Lrat-Cre*; NuTRAP mice across liver and adipose tissues.**

(A) Immunofluorescence staining of liver sections from *Lrat-Cre*; NuTRAP mice fed an HFCC diet, stained for GFP (green), αSMA (active hepatic stellate cell marker, red), and Hoechst (nuclei, blue). (B) Immunofluorescence staining of brown adipose tissue (BAT), inguinal white adipose tissue (iWAT), and epididymal white adipose tissue (eWAT) showing GFP (green), PLIN1 (adipocyte marker, red), and Hoechst (blue). (C) Immunofluorescence staining of BAT, iWAT, and eWAT showing GFP (green), PDGFRα (adipocyte progenitor marker, red), and Hoechst (blue). Arrows indicate colocalization between GFP and the respective marker proteins, and arrowheads indicate cells positive only for GFP. Scale bars: 30 μm (A-C).

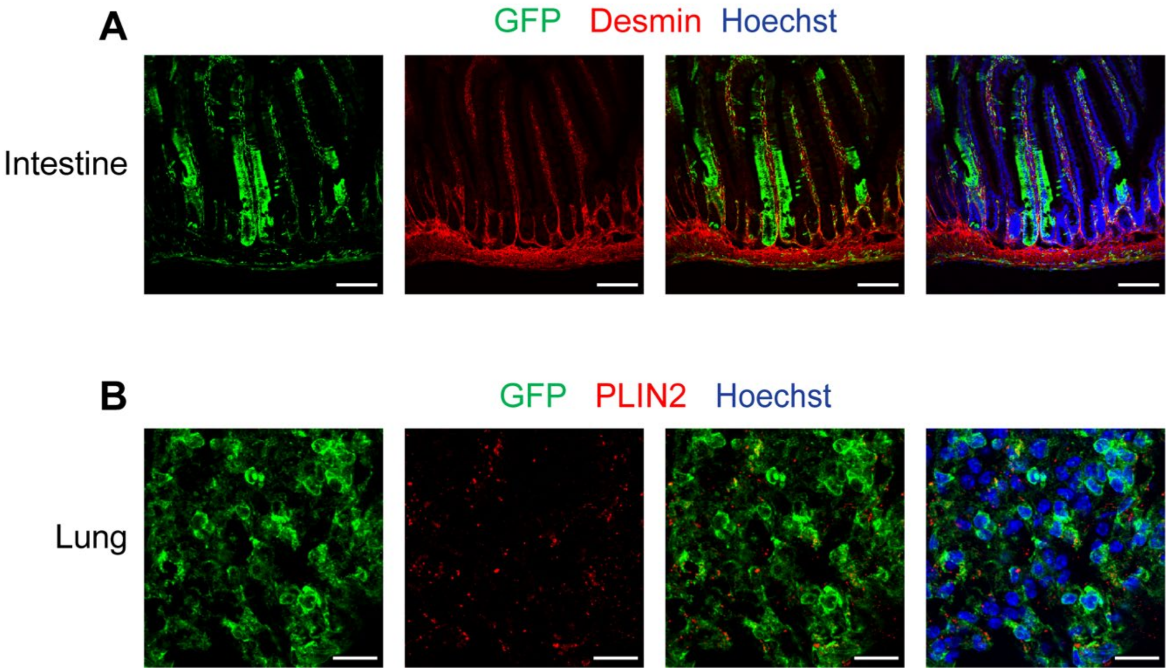


**Supplemental Figure 2. Distribution of GFP-positive cells in the intestine and lung of *Lrat-Cre*; NuTRAP mice**

(A) Immunofluorescence staining of intestinal sections stained for GFP (green), Desmin (smooth muscle marker, red), and Hoechst (blue) demonstrate robust GFP expression in enterocytes along the villi. Scale bars: 100 μm. (C) Immunofluorescence staining of lung sections stained for GFP (green), PLIN2 (lipofibroblast marker, red), and Hoechst (blue) reveals GFP expression in alveolar epithelial cells. Scale bars: 20 μm.

**SUPPLEMENTAL TABLE**

**Table S1**. Gene lists significantly enriched (1st tab) or depleted (2nd tab) in HSCs relative to whole liver. Genes were selected using the following cutoffs: average $\log_2$CPM ≥ 2, $\log_2$ fold change ($\log_2$FC) ≥ 1 and false discovery rate (FDR) < 0.05.
